# Intracranial recordings reveal ubiquitous in-phase and in-antiphase functional connectivity between homotopic brain regions in humans

**DOI:** 10.1101/2020.06.19.162065

**Authors:** Christian O’Reilly, Mayada Elsabbagh

## Abstract

Whether neuronal populations exhibit zero-lag (in-phase or in-antiphase) functional connectivity is a fundamental question when conceptualizing communication between cell assemblies. It also has profound implications on how we assess such interactions. Given that the brain is a delayed network due to the finite conduction velocity of the electrical impulses traveling across its fibers, the existence of long-distance zero-lag functional connectivity may be considered improbable. However, in this study, using human intracranial recordings we demonstrate that most interhemispheric connectivity between homotopic cerebral regions is zero-lagged and that this type of connectivity is ubiquitous. Volume conduction can be safely discarded as a confounding factor since it is known to drop almost completely within short inter-electrode distances (< 20 mm) in intracranial recordings. This finding should guide future electrophysiological connectivity studies and highlight the importance of considering the role of zero-lag connectivity in our understanding of communication between cell assemblies.

**Significance statement:** By analysing the functional connectivity between pairs of intracranial electrodes, we demonstrate that corresponding regions from the two cerebral hemispheres show oscillatory activity synchronized without any delay. Most often, these synchronized oscillations have an inverted amplitude, suggesting that the hemispheres inhibit one another in alternance. The existence of such synchronous activity between homotopic brain regions has important implications on how we understand communication between brain regions and on how we assess neuronal functional connectivity. Our results show that controlling the confounding effect of volume conduction by discarding instantaneous synchrony may result in missing important brain dynamics.

**Graphical abstract:** 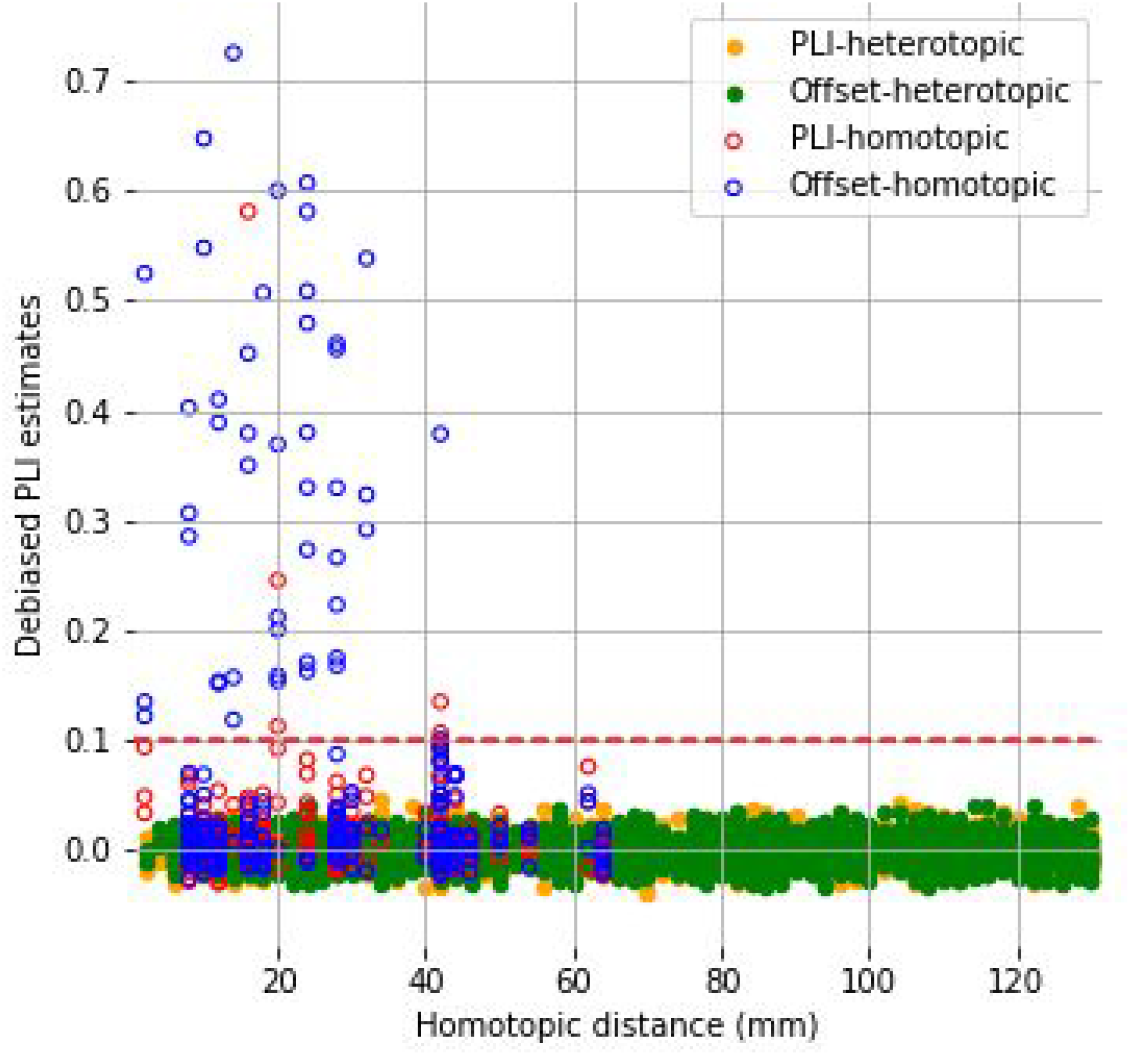

*Graphical abstract text:* Offset PLI and standard PLI (sensitive and insensitive to zero-lag connectivity, respectively) for pairs of electrodes separated by more than 105 mm, plotted against their homotopic distance. Significant connectivity (dots above the red dashed line) is found only between homotopic regions and mostly for zero-lag connectivity (offset-PLI).

## Introduction

The human brain is among the most complex networks known to humanity. It is estimated to comprise about 86 billion neurons (Azevedo et al., 2009), supported by as many glial cells (Azevedo et al., 2009; von Bartheld et al., 2016) and a sophisticated vasculature mesh. In the neocortex alone, neurons are interconnected by around 0.15 quadrillion synapses (Pakkenberg et al., 2003). Interactions within this vast network are enabled by mechanisms operating at distinct spatial and temporal scales. For example, while subcellular exchanges such as synaptic firing occur within milliseconds, other mechanisms like plasticity may involve days-long neuronal reorganization across broad cortical regions (Bassett and Sporns, 2017; Betzel and Bassett, 2017). Our understanding of these phenomena depends on our ability to measure brain activity across a spectrum of spatio-temporal scales.

The electroencephalogram (EEG) provides a time-resolved macroscopic correlate of cerebral activity. It can be recorded when synchronous discharges of spatially aligned populations of neurons, like the pyramidal cells, generate a cumulative electrical field that propagates by *volume conduction* from its source all the way to the scalp, where it can be picked up using surface electrodes. Unfortunately, the same volume conduction that makes EEG possible has been shown to be a major confounder for functional connectivity based on EEG (Nunez et al., 1997).

Unfolding this paradoxical role of volume conduction requires taking a closer look at the concept of EEG functional connectivity. Electrophysiological connectivity is generally considered a correlate of functional interaction due to saltatory conduction (i.e., the transmission of information through action potentials) propagated to single electrodes by short-range volume conduction. Ideally, there would be no long-range volume-conducted potentials reaching more than one electrode and no parallel means of communication, such as ephaptic coupling (Anastassiou et al., 2011) (i.e., modulation of neuronal computation by volume-conducted electrical fields). However, this ideal hangs on a very delicate trade-off between two opposite requirements; the neuronal activity must encompass a cell population sufficiently large to produce a measurable voltage deflection, but at the same time it must also be sufficiently spatially-restrained as to not overlap with regions covered by the other electrodes.

Moreover, the concept of functional connectivity, and the way it is reflected in EEG signals, shifts gradually across spatial scales. At a small scale, local connectivity is reflected directly in EEG amplitude since it depends on the synchronous activation of large neuronal populations. As we move to a larger scale, the connectivity can no longer be measured on a single EEG lead, but it rather relies on statistical dependence (e.g., coherence) between pairs of EEG signals generated by overlapping mixtures of neuronal sources. This overlap diminishes gradually as the scale — or equivalently the inter-electrode distance — becomes larger up to the point where volume conduction fully dissipates. In EEG, this distance is relatively large; for example, coherence measures have been determined to be significantly impacted by volume conduction for pairs of electrodes separated by at least up to 100 mm (Srinivasan et al., 1998). For smaller distances, an undetermined proportion of the statistical dependence is due to volume conduction from common neuronal sources rather than connectivity.

To overcome this severe limitation, neuronal communication based on action potentials needs to be distinguished from spurious dependencies due to volume conduction. Using the quasistatic approximation of Maxwell’s equations, volume conduction has been shown to propagate instantaneously for EEG (Plonsey and Heppner, 1967) and is therefore known to be a sufficient, <u>but not a necessary</u>, condition for in-phase or in-antiphase EEG activity between distant locations on the scalp. Such zero-lag synchronization has consequently often been downplayed as an artifact of volume conduction. For example, studies relying only on the *imaginary part of coherency* (Nolte et al., 2004) or the *phase lag index* (Stam et al., 2007), discard *by-design* any non-lagged activity between pairs of channels by considering only lagged synchronization. Alternatively, other approaches try to correct for the presence of linear leakage through orthogonalization before computing functional connectivity (Brookes et al., 2012; Colclough et al., 2015). In doing so, both approaches reject any zero-lagged synchronization contributing to the estimates of functional connectivity, regardless of whether it is an artifact or the result of genuine functional connectivity synchronized with no lag. This contrasts sharply with fundamental (Varela et al., 2001), experimental (Campo et al., 2019; Engel et al., 1991; Gray et al., 1989; Nikouline et al., 2001; Roelfsema et al., 1997), and modeling studies (Dalla Porta et al., 2019; Vicente et al., 2008; Viriyopase et al., 2012) supporting the existence of zero-lag neuronal connectivity.

Therefore, despite being treated as an artifact of volume conduction, zero-lag connectivity may be an important yet neglected mechanism underlying neural network communication. To investigate this possibility, we turned to intracranial recording (Frauscher et al., 2018), which is sensitive to volume-conducted activity from much smaller regions (< 20 mm (Bullock et al., 1995a, 1995b; Nunez et al., 1997)) than EEG (at least for < 100 mm (Srinivasan et al., 1998)) due to the use of smaller electrodes and the greater proximity between electrodes and neuronal sources. By introducing a one-sample offset between intracranial signals, we evaluated the proportion of zero-lag connectivity that is discarded from the standard PLI measure in electrodes distant enough (> 105 mm) not to share common volume-conducted sources. In doing so, we uncovered reliable and ubiquitous zero-lag connectivity – both in-phase and in-antiphase – between homotopic cortical regions.

## Materials and methods

### Intracranial recording dataset

For this study, we used an external open-access dataset of intracranial recordings (Frauscher et al., 2018) including 1772 channels recorded in 106 patients (54 males; mean age of 33.1 +/-10.8 years) who were candidates for surgical treatment of drug-resistant epilepsy. Included channels were from brain regions considered “very likely to be healthy” by a consensus of two epileptologists. Additional inclusion criteria included a sampling frequency of at least 200 Hz, the availability of peri-implantation neuroimaging allowing precise electrode localization, and the availability of control recordings performed at least 72h after implantation and at least 2h after any seizure. For each channel, 60 seconds of artifact-free recording was manually selected in an eyes-closed resting state, in rapid eye movement (REM) sleep, and in the second (N2) and third (N3) stages of non-REM sleep. The signals were filtered using a 0.5-80 Hz bandpass filter and downsample to 200 Hz when the recording sampling rate was higher. Electrode positions are provided in stereotaxic space. Corresponding brain regions for each electrode have been determined by coregistering a parcellation scheme consisting of 132 grey matter labels (Frauscher et al., 2018). These regions are used in the present study to determine homotopic pairs of channels. A more detailed description of inclusion criteria, signal preprocessing, and atlas co-registration is available in the original publication (Frauscher et al., 2018).

### Ethical approval

Data collection was performed under ethical approval granted to the Montreal Neurological Institute (REB vote: MUHC-15-950) as disclosed in the original paper (Frauscher et al., 2018).

### Connectivity assessment

To assess connectivity, the full 60 s segments are first split in 1 s epochs. Then, we computed the phase lag index (PLI) defined as follow (Ortiz et al., 2012; Stam et al., 2007):

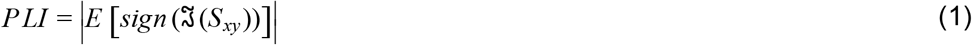

with *S_xy_* representing the cross-spectrum, 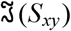 standing for the imaginary part of *S_xy_*, and E[x] denoting the expected value of x, which in practice is estimated by averaging across epochs. This measure assesses the degree to which the phases of two signals are locked in time. Importantly, it is defined such that all zero-lag activity (i.e., oscillatory activity with no phase difference) is explicitly canceled-out. Note that the zero-lag cancelation is not sensitive to differences in amplitude, and therefore, to amplitude inversion. This implies that phase lags of 180 degrees are also discarded from estimated connectivity. Mathematically, this comes from the fact that 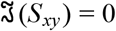 (i.e., it do not contribute to non-zero PLI estimate) when the angle of *S_xy_* is either zero degree (*S_xy_* reduces to a positive real number) or 180 degrees (*S_xy_* reduces to a negative real number). Consequently, we qualify as “zero-lag connectivity” any connectivity involving a phase difference that is an integer multiple of 180 degrees.

PLI is known to be a biased estimator of functional brain connectivity (Vinck et al., 2011). This is the case for any functional connectivity measure that is bound between 0.0 and 1.0 because their average value will consistently be greater than 0.0 on a random dataset. In our analyses, we estimated this bias by computing PLI on a surrogate dataset (surrogate-PLI) obtained by randomly selecting a subset of 200 channels and pairing every signal that came from different subjects. The mean value of this surrogate PLI has been removed from raw PLI estimates to obtain debiased PLI values.

We further compared the values of two PLI estimates, the standard (debiased) PLI measure and a second measure that we refer to as (debiased) offset-PLI. This second measure is computed exactly like the first one, with the only difference being that one of the two signals has been offset by a single time sample, allowing us to recover the zero-lag activity that was canceled-out by the first measure. Our intracranial signals being sampled at 200 Hz, this corresponds to a 5 ms offset. The average magnitude of the connectivity at this delay can be shown not to be significantly different from other similar delays (e.g., 10 or 15 ms; see Supplementary Figure 1.a). Consequently, the portion of the connectivity that is not recovered by the offset-PLI due to the cancelation of the 5 ms-lagged connectivity is relatively negligible. This contrasts with the very significant drop in the values of the offset-PLI when ignoring zero-lag correlations, as shown in Supplementary Figure 1.a by the very sharp and narrow drop in offset-PLI values at zero-lag. Although PLI is not linearly decomposable by time-delays, the difference between PLI and offset-PLI values can serve as a proxy for zero-lag connectivity in electrode pairs separated by distances larger than the range of volume conduction.

### Phase differences

The distribution of phase differences (Δ(ϕ)) between pairs of channels is determined by computing the angle from a polar representation of the cross-spectral density estimated using Welch’s averaged periodogram method (Welch, 1967). The *dominant peak phase discriminator* (ϕ_*d*_) indicating whether the dominant peak is at 0 or 180 degrees was evaluated using:

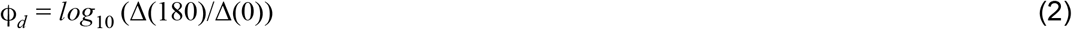

With this definition, a value of 1 indicates a phase distribution 10 times larger at a phase of 180 degrees than at 0 degrees, whereas a value of −1 indicates the opposite.

### Software

All analyses, including PLI computation, have been performed in Python 3.7.7 using MNE 0.20.dev0 and the standard Python libraries for signal processing, statistical analysis, and data visualization (Numpy 1.18.2, Scipy 1.4.1, Pandas 1.0.1, Matplotlib 3.1.2, Seaborn 0.9.0). Violin plot distributions have been generated using Seaborn with default parameters.

## Results

### Homologous brain regions exhibit ubiquitous zero-lag connectivity

We measured PLI and offset-PLI functional connectivity between every pair of electrodes in every subject. We also computed the surrogate-PLI to establish a chance level baseline. As shown in Figure 1.a, there is a bias of approximately 0.1 for PLI computed on 1-second epochs irrespective of the frequency. Thus, we debaised any further PLI estimates by subtracting mean surrogate-PLI value (mean ± sd: 0.102 ± 0.013; N=78,303).

**Figure 1.**
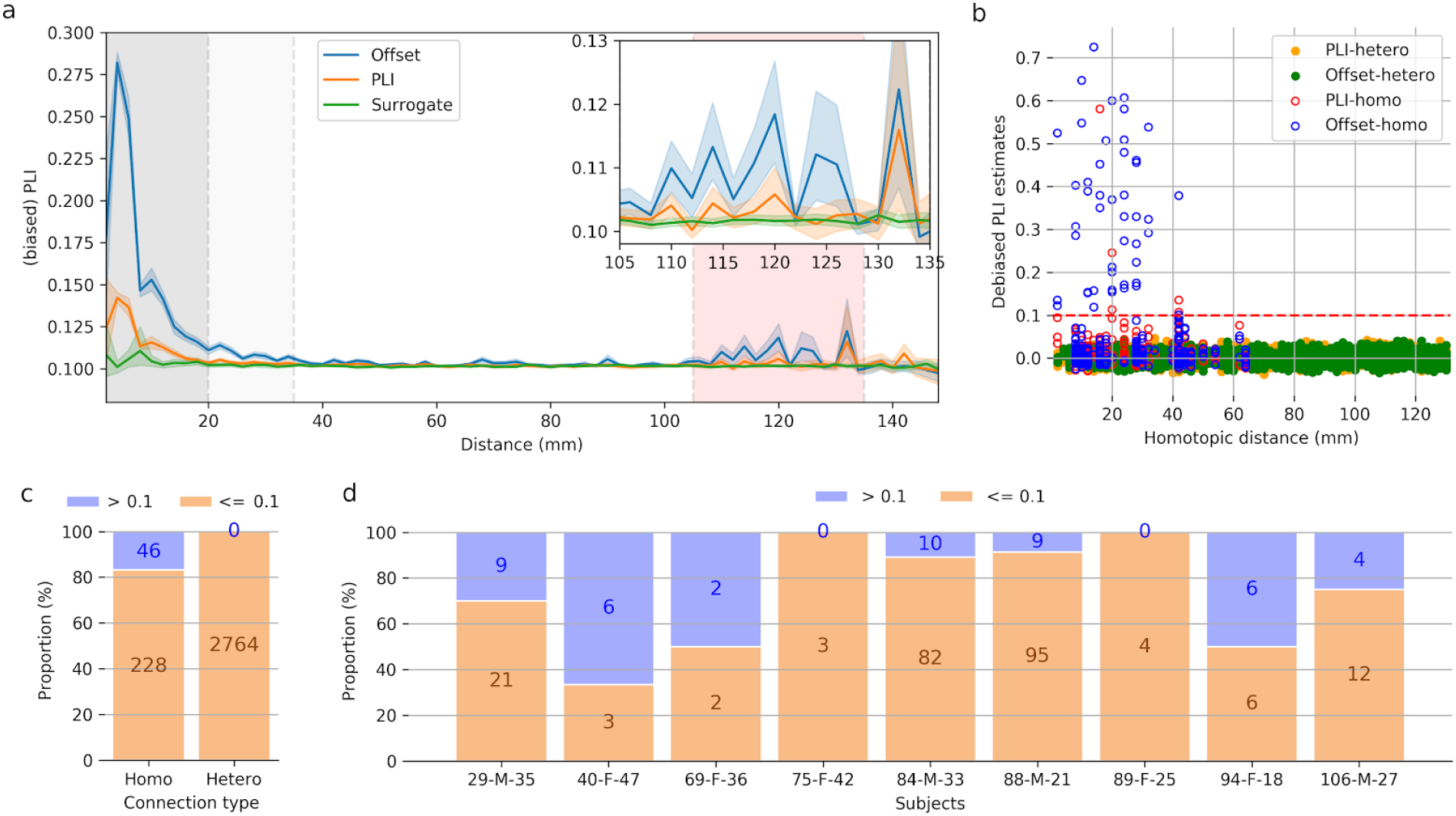
Standard and offset-PLI connectivity between pairs of channels. a) PLI values with respect to inter-electrode distances for the standard PLI definition (PLI; orange), the one-sample offset-PLI (offset; blue), and the PLI computed on surrogate data (surrogate; green). Shaded regions around the lines indicate the 95% confidence interval, computed using bootstrapping. The dark gray region delimits the range of distances for which we expect the presence of volume conduction. The pale gray region delineates a “safety margin” in which we do not interpret the connectivity results because of the possibility that some effect of volume conduction remains. The pink region delimits a range of distances for which significant connectivity is observed and for which we are confident that there cannot be any volume conduction artifact for intracranial recordings. We refer to the distances within this range as target distances since they are the focus of our subsequent analyses. The inset shows a close view of the PLI for the target distances. b) Offset (blue open circles: homotopic pairs; green filled circles: heterotopic pairs) and standard PLI (red open circles: homotopic pairs; orange filled circles: heterotopic pairs) for pairs of electrodes within the target distance range plotted against their homotopic distance (see text for the definition). The red dashed line shows the 0.1 threshold used to define significantly connected pairs. c) Proportion of significantly zero-lag connected pairs (debiased offset-PLI > 0.1) for homotopic and heterotopic pairs. d) Proportions of the homotopic pairs that are significantly zero-lag connected, per subjects. Labels with format XX-Y-ZZ on the x-axis refers to the identification numbers used for the subjects in the database (XX), the subjects biological sex (Y), and their age (ZZ).

Zero-lag connectivity for results within the dark grey region (inter-electrode distance < 20 mm) of Figure 1.a are likely to be heavily inflated by volume conduction. The offset-PLI is still significantly different from the surrogate-PLI estimates up to around 35 mm. Although at such distances volume conduction should not have a major effect, any difference between surrogate-PLI and offset-PLI values is debatable. In contrast, we observe offset-PLI significantly above surrogate-PLI values within an approximate range of 105 to 135 mm (shown as a pink area in Figure 1.a), which is clearly far beyond the reach of volume conduction in intracranial recordings. As opposed to offset-PLI, estimated standard PLI are barely larger than surrogate-PLI within this target range with confidence intervals generally overlapping with surrogate-PLI estimates (see inset in Figure 1.a), clearly demonstrating that most of the significant long-range connectivity is happening without any lag.

Having demonstrated significant zero-lag long-range connectivity, we further investigated the possibility that it may reflect interhemispheric connections between homotopics regions. We focused on within-subject electrode pairs separated by distances that fall within the 105 and 135 mm range. For these pairs, we computed their *homotopic distance*, defined as the distance between two channels if the left hemisphere was mirrored over the right hemisphere. In practice, since the mediolateral axis is centered on the interhemispheric fissure, this distance is obtained by using the absolute value of the channel coordinate along this axis.

We plotted the relationship between the homotopic distance and the standard PLI and the offset-PLI values, separately for homotopic and heterotopic channel pairs (Figure 1.b). It distinctly shows that all strong long-distance (> 105 mm) connections are established between channels with relatively short homotopic distances (≤ 42mm). To determine which pairs are considered significantly connected, we set a decision threshold on PLI values at 0.1. This criterion was chosen to separate adequately the sparse cloud of highly connected pairs from the dense cloud of lower values (see Figure 1.b). Using this threshold, within our target distance range, we found 46 significantly zero-lag connected channel pairs. From this subset, all pairs were connecting regions considered to be homotopic according to an atlas-based parcellation of the brain (see methods). By contrast, such *homotopic pairs* (i.e., channel pairs from homotopic regions) constitute only 7.6% (N=228/2992) of the pairs with significant offset-PLI values. Among all homotopic pairs, 16.8% (N=46/274) were showing significant debiased offset-PLI, whereas this ratio was of 0% (N=0/2764) for heterotopic pairs (Figure 1.c). The large number of heterotopic pairs (N=779/2764) separated by homotopic distances ≤ 42mm (i.e., the distance within which the significantly zero-lag connected pairs were found) indicates that it is the fact that two channels are implanted in homotopic regions and not that they are implanted at short homotopic distances that is the definitive factor for the existence of strong zero-lag connectivity, suggesting that the atlas-based segmentation of the brain accurately identifies functionally related regions. For the same electrode pairs, only six (all between homotopic regions) showed significant standard debiased PLI, confirming our previous conclusion that most of the interhemispheric connectivity is happening at zero-lag.

Highly connected pairs were observed in seven out of nine patients implanted with electrodes in homotopic regions (Figure 1.d). The two remaining subjects had very few homotopic pairs (3 and 4 pairs, respectively). This large proportion of homotopic pairs highly connected with zero-lag is surprising considering that they were obtained from only 60-second-long recordings of spontaneous activity, per subject and vigilance state. This observation indicates that zero-lag connectivity between homotopic regions is not a rare or exotic phenomenon occuring only in specific conditions, but a very common and ubiquitous mechanism.

Regarding our choice for the value for the significance threshold, it is worth mentioning that our conclusions are robust with respect to our choice for the significance threshold. A much smaller value could be used without affecting our conclusions since only homotopic pairs would be found for thresholds as low as 0.04. By choosing a comparatively high threshold (i.e., 0.10 instead of 0.04), we ensure that the pairs that enter subsequent analyses are strongly connected and that properties such as in-phase versus in-antiphase synchrony can be accurately estimated. Further, there is no apparent need to choose a more stringent threshold

(i.e., a higher value). If we consider the heterotopic pairs as representative of the distribution for the null hypothesis (i.e., not being zero-lag-connected beyond chance level), the p-value associated with pairs that have a 0.1 offset-PLI value is already vanishingly small (p-value=3.3e-16; one-tail z-test with μ=-8.063e-4 and σ=0.01246). Hence, this threshold is associated with a very strong statistical significance.

### Zero-lag connectivity often involves amplitude inversion

The first 2.5 seconds of activity is shown in Figure 2.a for three typical homotopic pairs with an offset-PLI > 0.1. The overlaid signals are tightly synchronized without any phase lag, but amplitude-inverted across the two hemispheres. Equivalently, we can say that these pairs of signals are synchronized in antiphase since amplitude inversion is mathematically equivalent to a 180-degree phase-shift for narrow-band oscillatory signals. Further, since PLI estimates account only for phase synchronization, irrespective of amplitude differences, any in-phase and amplitude inverted (eq. in-antiphase) synchronization between two signals is discarded by PLI estimates. Therefore, zero-lag connectivity uncovered by comparing offset-PLI with standard PLI estimates should be understood as connectivity between signals shifted by any integer multiple of 180 degrees, including zero degrees.

**Figure 2.**
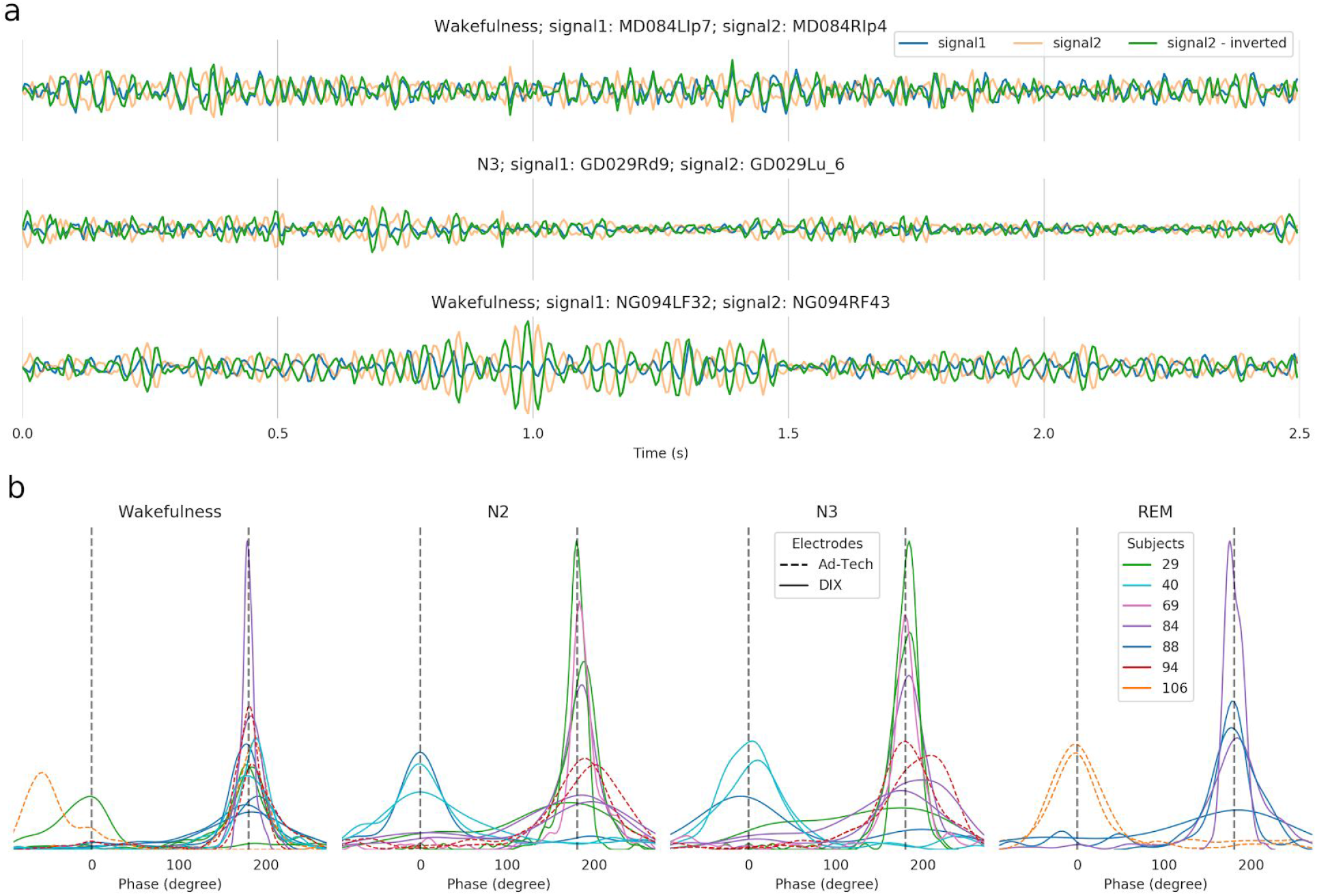
Amplitude inversion (eq. 180-degree phase-shift) in zero-lag connectivity. a) First 2.5 seconds of signals from three pairs of channels from homotopic regions that are highly connected with zero-lag connectivity. One of the signals is shown both with and without an amplitude inversion. The title above these three plots identifies the vigilance state and the database unique identifier of the two channels being plotted. b) Distribution of phase differences for all significantly zero-lag connected pairs. REM: Rapid eye movement sleep; N2 and N3: Second and third stages of non-REM sleep.

To illustrate the proportion of synchrony at zero (in-phase) versus 180 degrees (in-antiphase), we plotted the distribution of the phase difference between homotopic pairs with offset-PLI > 0.1 (Figure 2.b). The presence of well-defined peaks at either zero or 180 degrees for most pairs strongly confirms our findings of zero-lag connectivity. Although peaks can be observed at both phase-shifts, there are significantly more peaks (78%, Table 1) at 180 degrees vs. zero degrees (p = 2.0e-05; testing against the Bernoulli distribution under the null hypothesis that the dominant phase is equally likely to be at 0 or 180 degrees). Although this test is not exact given that the 46 electrode pairs include repeated measurements on subjects and channels, the dominance of the 180-degree offset appears robust since it is observed across all vigilance states and channel types. This was also the case in most subjects (5/7) showing significant zero-lag connectivity, where the percentages of 180-degree phase-shift dominance are 100%, 100%, 100%, 89%, 78%, 33%, and 25% respectively. Due to a large effect size (Cohen’s d=0.83; (Cohen, 1988), the dominance of the 180-degree phase-shift is statistically significant across subjects (t=2.03; p=0.044; one-tail one-sample t-test, compared against μ=50%) even with this small sample size (N=7). Further, using a surrogate approach (creating random samples with the same sizes as in our sample of significantly zero-lag connected homotopic pairs, but randomly pairing signals between-subjects to break any phase relationship, and bootstrapping with 10,000 iterations), we obtain a mean 180-degree phase-shift dominance of 50.0 ± 8.40 % (mean ± SD). When compared with the mean dominance across-subjects we obtained previously (75%), the surrogate distribution supports the presence of a 180-degree phase-shift dominance with p-value=0.0012 (one-tailed).

**Table 1.** The number of homotopic pairs with significant zero-lag connectivity that are either dominated by a 0 or a 180 degrees phase-shift, per vigilance state, and electrode type.

| State | Electrode | 0 degree | 180 degree | Percentage |
| --- | --- | --- | --- | --- |
| N2 | DIXI | 3 | 7 | 70% |
|  | Ad-Tech | 0 | 2 | 100% |
| N3 | DIXI | 3 | 7 | 70% |
|  | Ad-Tech | 0 | 2 | 100% |
| REM | DIXI | 0 | 5 | 100% |
|  | Ad-Tech | 2 | 0 | 0% |
| Wakefulness | DIXI | 1 | 10 | 91% |
|  | Ad-Tech | 1 | 3 | 75% |
| **Total** |  | **10** | **36** | **78%** |

### Zero-lag connectivity is independent of physiological and recording conditions

We examined the extent to which zero-lag connectivity is modulated by anatomical or physiological factors, namely, brain region (Figure 3.a,d), vigilance state (Figure 3.b), recording electrode type (Figure 3.c,e), and frequency band (Figure 3.f). Similar proportions were observed across brain lobes and vigilance states. Further, Kruskal-Wallis non-parametric one-way ANOVAs showed no significant effect of lobes or vigilance states on debiased offset-PLI values in homotopic pairs of electrodes. However, we found a significant effect of brain regions (H=29.1; p-values=5.9e-5). This is due only to pairs within the Planum Temporale. When data from this region are excluded, the region effect becomes non-significant. Most pairs in the Planum Temporal have a high offset-PLI (see Figure 3.d). However, detailed examination (see the full table of pairs with offset-PLI > 0.1 in Supplementary Table 1) shows that only two subjects are implanted bilaterally in the Planum Temporale and that the higher values come only from one of these subjects.

**Figure 3.**
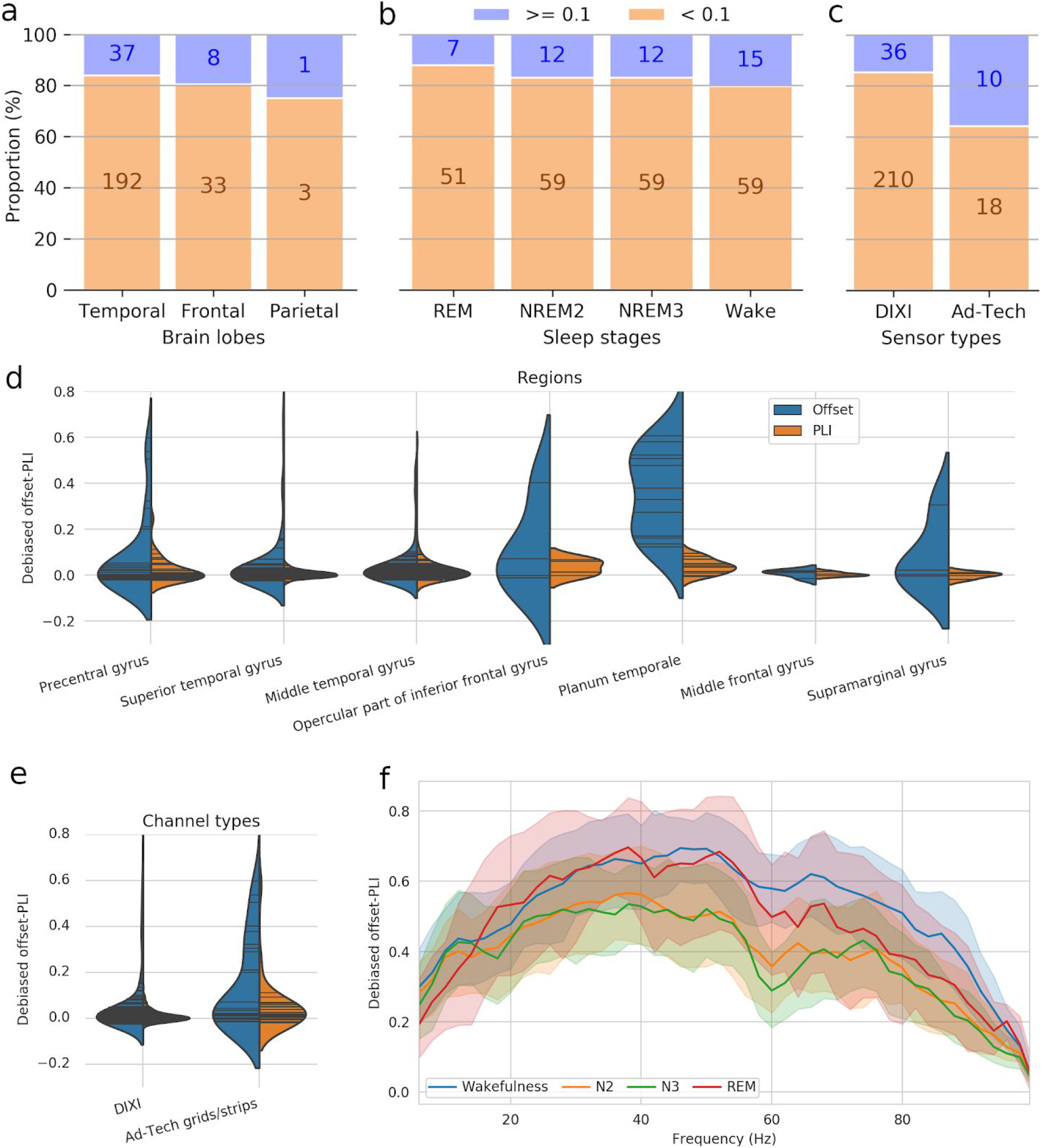
Modulation of zero-lag connectivity by physiological or experimental factors. a-c) Proportion of the homotopic pairs that show significant zero-lag connectivity, when splitting the sample per brain lobes (a), vigilance states (b), and sensor types (c). d,e) Kernel density estimation of the distribution of debiased PLI and offset-PLI values between homotopic channels for the different brain regions (d) and electrode types (e). Sticks show the individual data points. f) Variation of the debiased offset-PLI with frequency. PLI values are averaged, per vigilance state, across the significant homotopic pairs.

Regarding the electrode type, all homotopic pairs involved the same type of electrodes in both hemispheres, and these types were either DIXI electrodes (90%; 246/274) or Ad-Tech subdural strips and grids (10%; 28/274). We found a significant effect of the electrode type on offset-PLI values (H=6.52; p-values=0.011; see distributions in Figure 3.e). However, since both types of electrodes are well represented in the significantly zero-lag connected homotopic pairs, we can rule out the possibility that zero-lag interhemispheric connectivity is linked to a specific type of electrode.

Averaged offset-PLI computed per 2 Hz frequency bands for the set of channel pairs with debiased offset-PLI > 0.1 show that the zero-lag synchronization is broadband (see Figure 3.f; single curves, for all 46 significant pairs, are also provided in Supplementary Figure 2), reflecting that large populations of neurons can fire synchronously across brain regions following non-sinusoidal patterns.

## Discussion

### Potential roles of zero-lag connectivity

Using intracranial recordings, we demonstrated ubiquitous zero-lag connectivity between homotopic brain regions, which was three times more often in antiphase than in phase. It is well known that the polarity of local field potentials and current source densities can alternate depending on the laminar depth within a given cortical column (Schaefer et al., 2015). Thus, it is possible that some amplitude inversions we observed are due to electrodes recording in homotopic regions at different cortical depths. Nevertheless, this is unlikely to explain the relatively high and robust predominance of in-antiphase synchronization observed in our data.

In general, neocortical neurons tend to synchronize with small but reliable delays, which have been shown to encode stimulus properties (Uhlhaas et al., 2009). Since our results indicate clear and reliable zero-lag synchronization, this homotopic connectivity is likely to serve a different function. For example, a mechanism of competition/reciprocal-inhibition is probably involved in the observed antiphase synchronization. Previous studies using transcranial magnetic stimulation have identified a similar pattern of interhemispheric inhibition over the primary motor area (Ferbert et al., 1992; Perez and Cohen, 2009). However, to our knowledge, this is the first time phase-synchrony is reported for spontaneous interhemispheric inhibition and the fact that this synchrony involves no lag is an outstanding observation.

Regarding in-phase connectivity, synchronized oscillatory activity between two brain regions, resulting from generalized depolarization (excitation) and hyperpolarization (inhibition) at the population level can reasonably be expected to creates common windows of heightened and reduced responsiveness, respectively. Therefore, it can be hypothesized that such a window promotes information integration between these distant regions as formulated by the *Communication through Coherence* hypothesis (Fries, 2015). Such in-phase synchrony could be hypothesized in some cases to come from common sub-cortical drivers projecting to both hemispheres (Huang and Pipa, 2007). For example, this would be the case for thalamocortical rhythms (Steriade et al., 1993). However, there is also evidence for long-distance synchronization to be mediated through reciprocal cortico-cortical projections (Uhlhaas et al., 2009). Further, reciprocal cortico-thalamo-cortical projections could also support the emergence (i.e., not imposed by a sub-cortical driver) of zero-lag synchronization (Vicente et al., 2008).

Zero-lag connectivity may also be involved in supporting the brain’s predictive capacity. The brain is a delayed network due to the finite conduction velocity of the electrical impulses traveling across its fibers. In such a network, the apparent paradox of zero-lag (instantaneous) connectivity can be resolved through rhythmic entrainment supported by feedback in closed-loop systems. By capturing the rhythm by which a phenomenon of interest occurs (i.e., rate coding), neuronal populations can synchronize their activity with expected external or internal cues, allowing anticipation and better response time. Such synchrony would need to occur bilaterally for any process requiring operations performed either in both hemispheres at the same time, or potentially in either hemisphere, depending on the context (e.g., the laterality of a stimulus).

The fact that we did not find a clear modulation of the zero-lag connectivity by the vigilance state may come as a surprise given that arousal levels have a strong impact on scalp EEG synchrony. This may be due partly to the fact that the EEG and intracranial recordings are sampling the neuronal population at different spatial scales. Neuronal activity that is strongly synchronized in a very local network may register on implanted electrodes but not on scalp EEG, whereas weaker but spatially extended neuronal synchrony may register on EEG but not on intracranial electrodes. Further, strong synchronization associated for example with deep sleep slow waves is known to present as traveling waves (Botella-Soler et al., 2012; Massimini et al., 2004) and, therefore, involves a propagation delay and does not contribute to long-distance zero-lag connectivity.

### Implications for functional connectivity in EEG

Our findings of ubiquitous zero-lag connectivity between homotopic brain regions have important implications for the measurement of functional connectivity using EEG. Traditional solutions to deal with the confounding effects of volume conduction include studying only long-range connections (e.g., considering only pairs of electrodes separated by more than 10 cm in a coherence study; Srinivasan et al., 1998) or discarding any zero-lag connectivity (e.g., using the imaginary part of coherency or the PLI; Nolte et al., 2004; Stam et al., 2007). In contrast, our findings clearly demonstrate the importance of zero-lag connectivity, indicating that the confounding effect of volume conduction needs to be controlled for without systematically discarding zero-lag connectivity. Instead, functional connectivity studies using EEG should adopt strategies to un-mix cortical sources such as independent component analysis (Delorme et al., 2012) or source reconstruction (Baillet et al., 2001), or use model-based approaches such as dynamic causal modeling (Kiebel et al., 2009). The latter approach may be argued to be more powerful; with adequate modeling, it could take into account other factors that are currently neglected, such as ephaptic coupling or long-distance neurochemical mediation. A model-based approach can also be used to inject more a priori knowledge or benefit from integrating data from other modality, as demonstrated with estimates of functional connectivity informed by structural connectivity (Glomb et al., 2020). It should nevertheless be kept in mind that un-mixing strategies are trying to solve a massively underdetermined problem using heuristic and only partially succeed in doing so, leaving spurious zero-lag synchrony between sources (i.e., a problem known as source leakage) that, much like volume conduction, needs to be controlled for. Hopefully, with future advances in inverse modeling, the degree of source leakage will be reduced. In the meantime, researchers not wanting to throw the baby with the bathwater should probably develop the habits of testing and comparing the outcome of at least two metrics of connectivity in their study, one that discards zero-lag synchrony (offers robustness against volume conduction and source leakage) and one that does not (is sensitive to genuine zero-lag connectivity). Although the thorny problem of distinguishing what part of the zero-lag synchrony is genuine and what part constitutes an artifact is still an open issue, we argue that it would be misguided to discard the problem altogether by ignoring the clear existence of genuine zero-lag connectivity.

## Limitations

The study of human intracranial recordings comes with challenges inextricably linked to the fact that the choice of the participants and of the electrode placement for these recordings is strictly dictated by medical imperative. Hence, our sample of subjects with homotopic recordings is relatively small and it comes only from epileptic patients. Although steps have been taken to ensure that the recordings used here are from healthy brain regions (see methods), it is not possible to totally exclude the possibility of biases being introduced due to the ethical constraints set on the selection of the participants and the implanted brain regions. These results should be considered alongside other modes of investigations to gain a more complete understanding of zero-lag connectivity in the brain.

Further, the impact of demographic variables such as age and sex could not be studied in this study given that the small number of subjects implanted with electrodes in homotopic regions (N=9) would not provide a sufficient statistical power.

## Conclusion

In this study, we found ubiquitous in-phase and in-antiphase connectivity between homotopic regions of the brain. This zero-lag activity was highly specific to homotopic pairs of channels, which rules out any possibility that this observation is due the method confounders such as volume conductions spreading on unexpectedly long distances or correlations induced by improper use of reference electrodes since any of these factors would affect equally homotopic and heterotopic regions. Further, since it is impossible for a single source to create spatially discontinuous inphase patterns (Tognoli and Kelso, 2009), the observation of inphase synchrony between long-distance homotopic regions and not between smaller-distance heterotopic regions is unlikely to be due to volume conduction.

These results have important implications in our understanding of functional connectivity. Our findings demonstrate the preponderance of zero-lag connectivity and suggest that it probably serves fundamental mechanisms relevant to the integration and interpretation of input across distinct brain regions. Since this type of connectivity has been found specifically between homotopic regions, it may be particularly impacted by conditions altering the structural connectivity in the corpus callosum, like in the case of autism spectrum disorder for example (Travers et al., 2015). In light of these results, it is clear that such synchronous activity should not be excluded from EEG connectivity analyses as an artifact.

## Conflict of Interest Statement

Authors report having no conflict of interests.

## Author Contributions

C.O.R.: Conceptualization, data analysis, writing of the first manuscript draft, revision of the manuscript.

M.E.: Revision of the manuscript.

## Acknowledgments

This research is supported by the Azrieli Centre for Autism Research (ACAR). The authors would like to thank Samantha Wunderlich for proofreading this manuscript.

## Data and code availability

The intracranial EEG database is available at https://mni-open-ieegatlas.research.mcgill.ca. The code used for the analysis is accessible on GitHub (https://github.com/christian-oreilly/intracranial_paper).

## Supporting information

### Effect of delays and epoch durations on offset-PLI

Supplementary Figure 1 shows averaged offset-PLI values for the four different vigilance states at various delays. To ensure that interpretation of these figures is not confounded by volume conduction, they have been produced using only pairs of electrodes separated by at least 35 mm and we included only pairs for which the overall debiased offset-PLI was larger than 0.1. Each time step corresponds to a 5 ms offset, given the recording sample rate (200 Hz). Note that the values at step=0 correspond by definition to the standard PLI. Panels a and b show offset-PLI computed using 1 s and 5 s epochs, respectively. The same scale has been used for these two panels to highlight the effect of epoch duration on PLI computation. Not only the bias is very much impacted (not visible on this figure using debiased offset-PLI; 0.225 ± 0.014 (mean ± sd) for 5s epochs as compared to 0.102 ± 0.013 for 1s epochs), but even debiased offset-PLI takes very different values. The average values of offset-PLI vary smoothly with respect to the number of time steps, except for step=0. On the one hand, the smoothness for steps different from zero supports that most connectivity is preserved when computing PLI at an arbitrary offset close to zero. On the other hand, the sharp drop at step=0 shows the importance of zero-lag connectivity, even for channel pairs that are not expected to be sensitive to volume conduction.

PLI is a biased estimator (Vinck et al., 2011), as can be clearly understood by the fact that its support is [0, 1]. This implies that any source of noise (i.e., in this case, any fortuitous and momentary signal cross-correlation) will systematically cause values greater than zero such that the estimated averages will also be larger than 0. The relationship between this bias and the epoch length has an impact on the absolute values of the (biased) PLI values, but it has no consequence on our results since 1) we debiased our PLI estimates, and 2) we always use the same epoch length throughout our analyses (except for our analysis of the impact of epoch duration).

### Explanation of the effect of the epoch duration

We observed that the duration of the epochs has a significant difference on the rate at which the offset-PLI values decrease with increasing offsets (Supplementary Figure 1.a,b). In order to explain this relationship, we propose a model where we first penalize the connectivity by a factor ε equal to the ratio of the offset to the epoch duration:

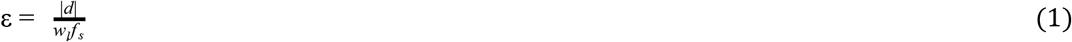

where *d* and *w_I_*, are the offset and the epoch length in sample numbers and *f_s_* is the sampling frequency. This penalty accounts for reduced bleeding of the offset-PLI zero-lag cancellation within neighboring frequencies in larger epochs due to a consequent increase in spectral resolution. Panels (d) and (e) of Supplementary Figure 1 show that the pattern of canceled frequencies at different delays remains the same for these two epoch durations, but for shorter epochs, it is more blended and smoothed. Panel (f) overlay the variation of PLI by frequencies for different offsets, whereas panel (g) enlarges a small segment of these curves for the case with an offset of 20 time samples (100 ms). The sharpness of the troughs at *frequencies where PLI discards zero-lag connectivity*^1^ (shown by vertical dashed lines) is clear for the 5 s epochs. By contrast, for the 1 s epochs, the zero-lag connectivity cancelation at two neighboring frequencies (e.g., 20 and 25 Hz on panel (g)) bleeds onto larger frequency intervals resulting in a very sharp and more heavily attenuated intermediate peak (e.g., at around 22.5 Hz on panel (g)). For larger epochs, the resulting higher frequency sampling is associated with smaller frequency intervals over which connectivity is canceled-out, resulting in a slower decrease of the offset-PLI as the offset increases.

While this increased attenuation due to reduced spectral resolution may account for most of the differences observed for offset-PLI across different epoch sizes, it does not account for the curved aspects of these relationships. Thus, we further scaled the ε penalty by a weight *w*(|*d*|) proportional to the square root of the area under the *1/f*^β^ aperiodic spectral component, normalized to unity and integrated from the inverse of the offset up to half the sampling frequency (i.e., the Nyquist frequency):

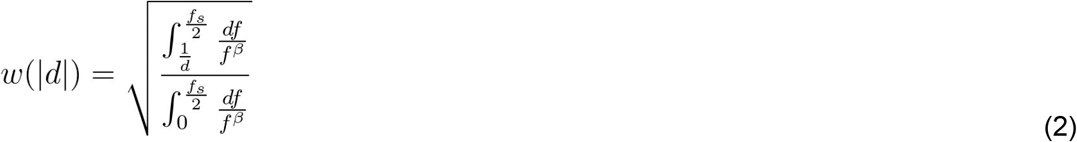

The square root in (2) is used to account for the fact that the *1/f*^β^ equation describes the aperiodic component of the *power* spectrum, which is related to the amplitude by a square relationship. β was set to 2.29, as previously reported (Frauscher et al., 2018). The *w*(|*d*|) factor accounts for the fact that higher frequencies are expected to be more severely affected by smaller delays because these delays constitute a larger fraction of their period. As the offset increases, we penalize the connectivity more because the offset starts to increasingly affect the lower frequencies while continuing to affect high frequencies. Finally, we scale the penalty by a single proportionality constant *a* fitted manually to a value of 12 in our analysis. The final relationship is as follow:

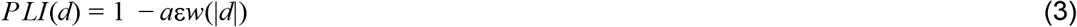

Supplementary Figure 1.c shows the variation of offset-PLI with respect to the offset as modeled with the equation (3), overlaid to the empirical curves shown in panel (a) and (b), but averaged across states and normalized to their peak. This tentative explanation captures relatively well the overall behavior displayed by the offset-PLI computed using different epoch durations, using only one free parameter (*a*).

Importantly, although we think this analysis was important to validate the behavior of the offset-PLI, it should be noted that these factors are not expected to have any impact on our main analyses because they indicate systematic differences in offset-PLI with respect to either the epoch duration or the value of the offset, which are both kept constant in our analysis across all offset-PLI estimates. Thus, any systematic bias would contribute equally to the mean value of the compared distribution and would therefore not induce between-condition (e.g., homotopic versus heterotopic) mean differences.

**Supplementary Table 1.**
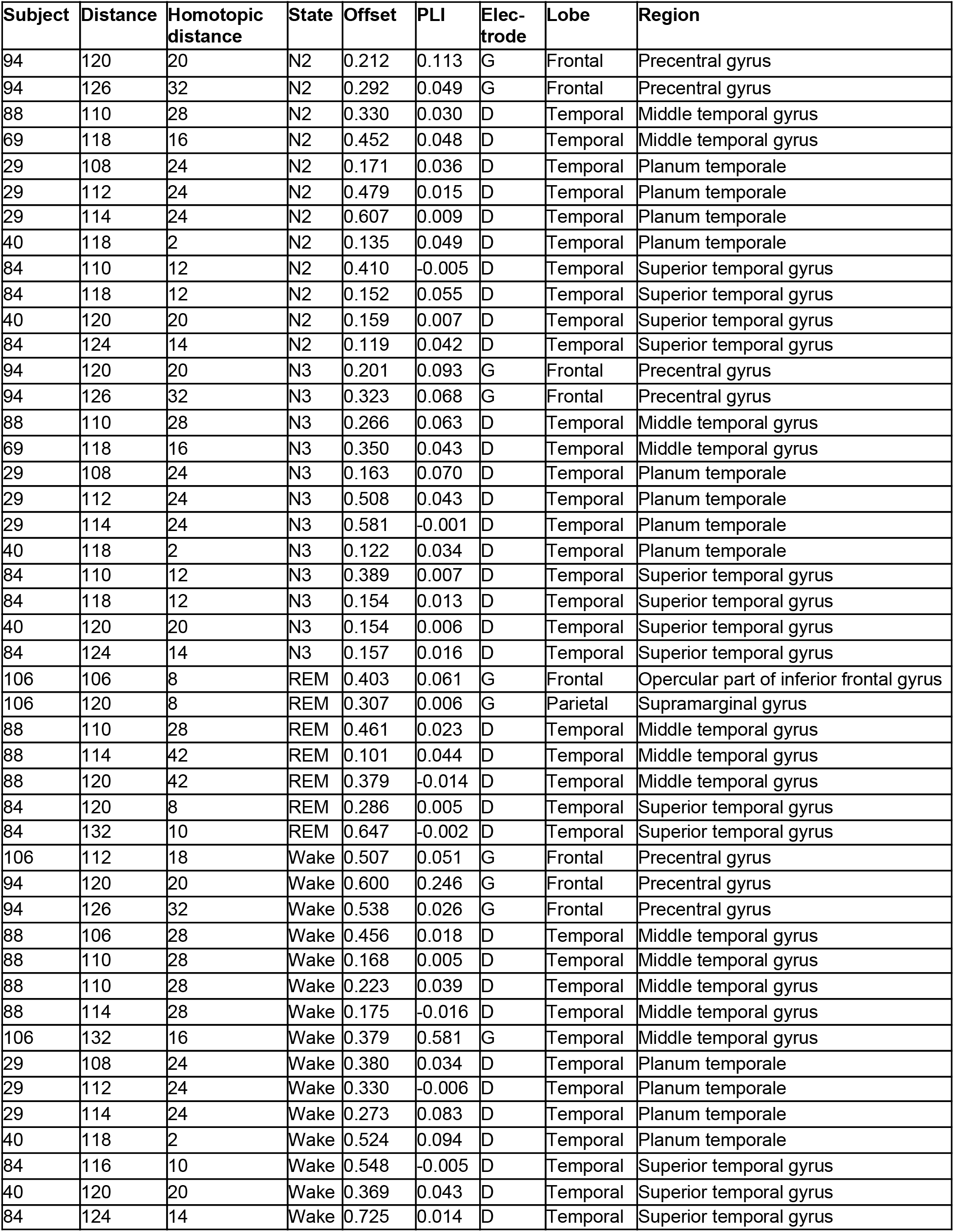
Significantly zero-lag connected (i.e., offset-PLI > 0.1) homotopic pairs.

| Subject | Distance | Homotopic distance | State | Offset | PLI | Elec-trode | Lobe | Region |
| --- | --- | --- | --- | --- | --- | --- | --- | --- |
| 94 | 120 | 20 | N2 | 0.212 | 0.113 | G | Frontal | Precentral gyrus |
| 94 | 126 | 32 | N2 | 0.292 | 0.049 | G | Frontal | Precentral gyrus |
| 88 | 110 | 28 | N2 | 0.330 | 0.030 | D | Temporal | Middle temporal gyrus |
| 69 | 118 | 16 | N2 | 0.452 | 0.048 | D | Temporal | Middle temporal gyrus |
| 29 | 108 | 24 | N2 | 0.171 | 0.036 | D | Temporal | Planum temporale |
| 29 | 112 | 24 | N2 | 0.479 | 0.015 | D | Temporal | Planum temporale |
| 29 | 114 | 24 | N2 | 0.607 | 0.009 | D | Temporal | Planum temporale |
| 40 | 118 | 2 | N2 | 0.135 | 0.049 | D | Temporal | Planum temporale |
| 84 | 110 | 12 | N2 | 0.410 | -0.005 | D | Temporal | Superior temporal gyrus |
| 84 | 118 | 12 | N2 | 0.152 | 0.055 | D | Temporal | Superior temporal gyrus |
| 40 | 120 | 20 | N2 | 0.159 | 0.007 | D | Temporal | Superior temporal gyrus |
| 84 | 124 | 14 | N2 | 0.119 | 0.042 | D | Temporal | Superior temporal gyrus |
| 94 | 120 | 20 | N3 | 0.201 | 0.093 | G | Frontal | Precentral gyrus |
| 94 | 126 | 32 | N3 | 0.323 | 0.068 | G | Frontal | Precentral gyrus |
| 88 | 110 | 28 | N3 | 0.266 | 0.063 | D | Temporal | Middle temporal gyrus |
| 69 | 118 | 16 | N3 | 0.350 | 0.043 | D | Temporal | Middle temporal gyrus |
| 29 | 108 | 24 | N3 | 0.163 | 0.070 | D | Temporal | Planum temporale |
| 29 | 112 | 24 | N3 | 0.508 | 0.043 | D | Temporal | Planum temporale |
| 29 | 114 | 24 | N3 | 0.581 | -0.001 | D | Temporal | Planum temporale |
| 40 | 118 | 2 | N3 | 0.122 | 0.034 | D | Temporal | Planum temporale |
| 84 | 110 | 12 | N3 | 0.389 | 0.007 | D | Temporal | Superior temporal gyrus |
| 84 | 118 | 12 | N3 | 0.154 | 0.013 | D | Temporal | Superior temporal gyrus |
| 40 | 120 | 20 | N3 | 0.154 | 0.006 | D | Temporal | Superior temporal gyrus |
| 84 | 124 | 14 | N3 | 0.157 | 0.016 | D | Temporal | Superior temporal gyrus |
| 106 | 106 | 8 | REM | 0.403 | 0.061 | G | Frontal | Opercular part of inferior frontal gyrus |
| 106 | 120 | 8 | REM | 0.307 | 0.006 | G | Parietal | Supramarginal gyrus |
| 88 | 110 | 28 | REM | 0.461 | 0.023 | D | Temporal | Middle temporal gyrus |
| 88 | 114 | 42 | REM | 0.101 | 0.044 | D | Temporal | Middle temporal gyrus |
| 88 | 120 | 42 | REM | 0.379 | -0.014 | D | Temporal | Middle temporal gyrus |
| 84 | 120 | 8 | REM | 0.286 | 0.005 | D | Temporal | Superior temporal gyrus |
| 84 | 132 | 10 | REM | 0.647 | -0.002 | D | Temporal | Superior temporal gyrus |
| 106 | 112 | 18 | Wake | 0.507 | 0.051 | G | Frontal | Precentral gyrus |
| 94 | 120 | 20 | Wake | 0.600 | 0.246 | G | Frontal | Precentral gyrus |
| 94 | 126 | 32 | Wake | 0.538 | 0.026 | G | Frontal | Precentral gyrus |
| 88 | 106 | 28 | Wake | 0.456 | 0.018 | D | Temporal | Middle temporal gyrus |
| 88 | 110 | 28 | Wake | 0.168 | 0.005 | D | Temporal | Middle temporal gyrus |
| 88 | 110 | 28 | Wake | 0.223 | 0.039 | D | Temporal | Middle temporal gyrus |
| 88 | 114 | 28 | Wake | 0.175 | -0.016 | D | Temporal | Middle temporal gyrus |
| 106 | 132 | 16 | Wake | 0.379 | 0.581 | G | Temporal | Middle temporal gyrus |
| 29 | 108 | 24 | Wake | 0.380 | 0.034 | D | Temporal | Planum temporale |
| 29 | 112 | 24 | Wake | 0.330 | -0.006 | D | Temporal | Planum temporale |
| 29 | 114 | 24 | Wake | 0.273 | 0.083 | D | Temporal | Planum temporale |
| 40 | 118 | 2 | Wake | 0.524 | 0.094 | D | Temporal | Planum temporale |
| 84 | 116 | 10 | Wake | 0.548 | -0.005 | D | Temporal | Superior temporal gyrus |
| 40 | 120 | 20 | Wake | 0.369 | 0.043 | D | Temporal | Superior temporal gyrus |
| 84 | 124 | 14 | Wake | 0.725 | 0.014 | D | Temporal | Superior temporal gyrus |

**Supplementary Figure 1.**
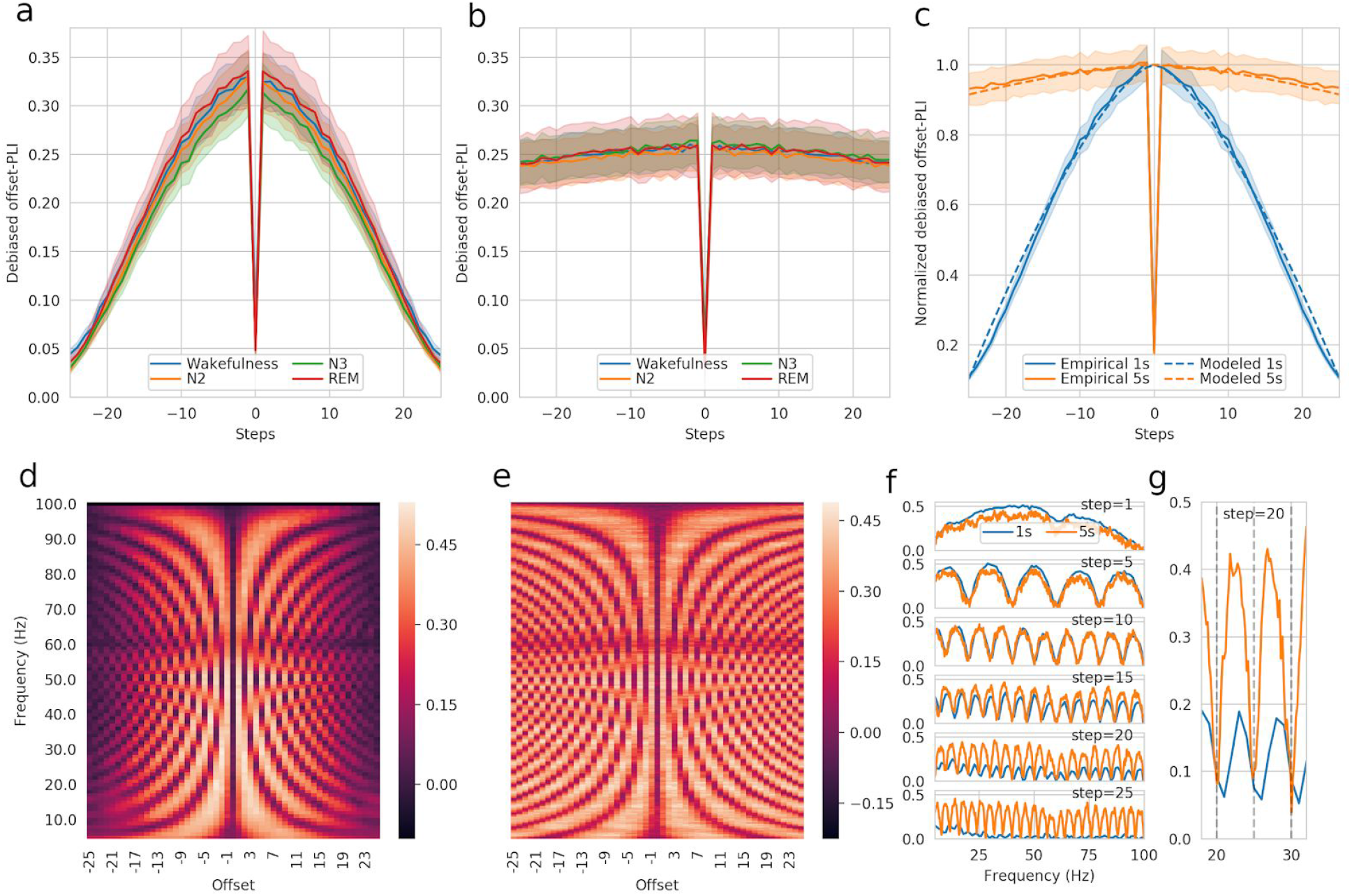
Impact of delays and epoch duration on PLI estimates. a, b) Variation of the offset-PLI for different offsets, for 1 s epochs (a) and 5 s epochs (b). c) Curves from panels (a) and (b) averaged across states and normalized to unity at their peaks. The relationships modeled using equation (3) have also been overlaid to these curves. d, e) Heat maps of offset-PLI values for a range of frequencies and offsets, computed using 1 s epochs (d) and 5 s epochs (e). f) Variation of the offset-PLI with respect to frequency, at discrete steps (i.e., number of time samples used as offset), overlaid for the two epoch lengths. g) A zoomed-in view of the 18-32 Hz range for the offset-PLI at an offset of 20 time samples (i.e., 100 ms).

**Supplementary Figure 2.**
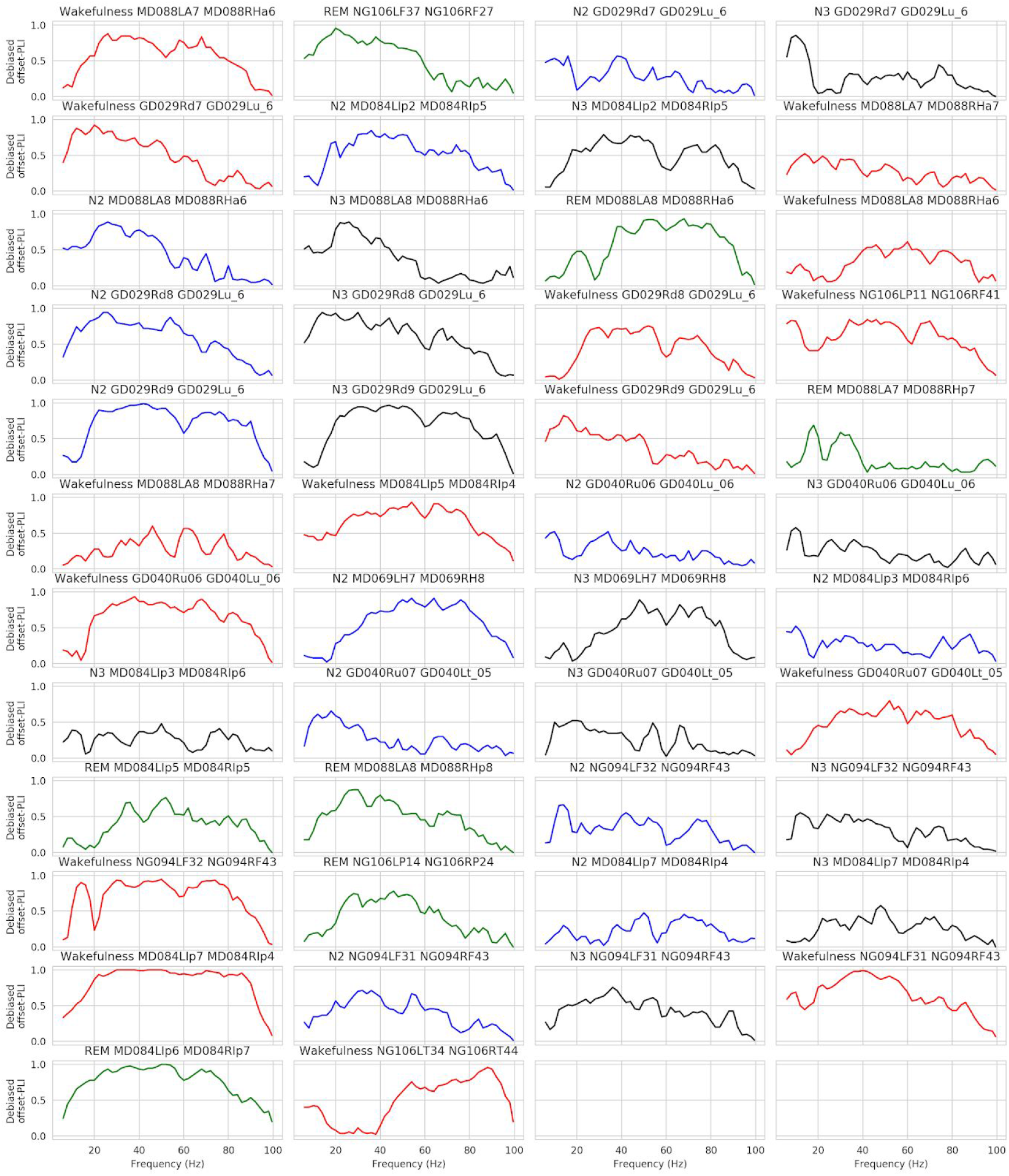
Variation of the debiased offset-PLI with respect to frequency for the 46 homotopic pairs with a global debiased offset-PLI greater 0.1. The title over the plots identifies the vigilance state and the database unique identifiers for the two channels. The vigilance state is color-coded: red for wakefulness, blue for N2, black for N3, and green for REM. The offset-PLI values have been computed on every contiguous 2 Hz frequency band.

## Footnotes

1 For a given offset Δ, the expression *“frequencies where PLI discards zero-lag connectivity”* can be understood as frequencies where the zero-lag connectivity is canceled-out both at offset=0 and offset=Δ. For example, with an offset of two time steps (a period of 10 ms), the activity that was zero-lagged at 50 Hz with offset=0 ms is again zero-lagged with offset=10 ms since the signal has been shifted by exactly half a cycle (i.e., 180 degrees) from its original phase.

## Notes

**Supporting or grand information**: This research is supported by the Azrieli Centre for Autism Research (ACAR).

### Competing Interest Statement

The authors have declared no competing interest.

### Summary of Updates

The discussion has been extended. Various modifications have been prompted by the peer-review process to clarify the text. The reported results remain unchanged with respect to the previous version.

